# Hyperspectral imaging has a limited ability to remotely sense the onset of beech bark disease

**DOI:** 10.1101/2024.09.20.614150

**Authors:** Guillaume Tougas, Christine I. B. Wallis, Etienne Laliberté, Mark Vellend

**Affiliations:** Université de Sherbrooke, Université de Montréal; Université de Sherbrooke, Technische Universität Berlin; Université de Montréal; Université de Sherbrooke

**Keywords:** Remote Sensing, Beech Bark Disease, Forest Pathogens, Hyperspectral Imaging, Microdrone

## Abstract

Insect and pathogen outbreaks have a major impact on northern forest ecosystems. Even for pathogens that have been present in a region for decades, such as beech bark disease (BBD), new waves of mortality are expected in host populations. Hence, there is a need for innovative approaches to monitor their advancement extensively in real-time. Here we test whether airborne hyperspectral imaging – involving data from 344 wavelengths in the visible, near infrared (NIR) and short-wave infrared (SWIR) – can be used to assess beech bark disease severity in southern Quebec, Canada. Field data on disease severity were linked to the airborne hyperspectral data for individual beech crowns. Partial least-squares regression (PLSR) models using airborne imaging spectroscopy data predicted a small proportion of the variance in beech bark disease severity: the best model had an R^2^ of only 0.10. Wavelengths with the strongest contributions were from the NIR (∼719 nm) and the SWIR (∼1287 nm), which may suggest mediation by canopy greenness, water content and canopy architecture. Similar models using hyperspectral data taken directly on individual leaves had no explanatory power (R^2^ = 0). In addition, airborne and leaf-level hyperspectral datasets were uncorrelated. The failure of leaf-level models suggests that canopy structure was likely responsible for the limited predictive ability of the airborne model. Somewhat better performance in predicting disease severity was found using common band ratios for canopy greenness assessment (the Green Normalized Difference Vegetation Index, gNDVI, the Red-edge Inflexion Point, REIP, and the Normalized Phaeophytinization Index, NPQI); these variables explained up to 19% of the variation in disease severity. Overall, we argue that the complexity of hyperspectral data is not necessary for assessing BBD spread and that spectral data in general may not provide an efficient means of improving BBD monitoring on a larger scale.

## 1. Introduction

Northern forests are becoming increasingly susceptible to insect outbreaks, largely due to a higher frequency of extreme weather events and rising temperatures (Harvey et al., 2020; Jactel et al., 2019; Johnson et al., 2010; Pureswaran et al., 2018; Volney & Fleming, 2000). These rapid changes can cause a decrease in net ecosystem productivity, prompting carbon sinks to become sources (Kurz et al., 2008). There is thus a major need for better methods of monitoring disease progression, both in terms of frequency and severity (Senf et al., 2017).

Throughout most of its native range, American beech (*Fagus grandifolia*) has been impacted by beech bark disease (BBD), a vascular disease attributed to *Neonectria* spp. fungi. Beech is a late-successional species in eastern North American hardwood forests, typically co-occurring with maple species. In long-affected regions, including most of the maritime provinces of Canada, southern Quebec and the northeastern United States, disease-free beech stands are almost non-existent (<1%, Cale et al., 2017; Garnas et al., 2013; Shigo, 1972). The disease arrived in North America with the introduction of the beech scale insect (*Cryptococcus fagisuga*) in Halifax, Nova Scotia, Canada, around 1890 (Ehrlich, 1934). Microlesions in the bark caused by the scale insect permit entry of *Neonectria* fungi, which are native to North America and historically only infected wounded beech trees (Ehrlich, 1934; Hirooka et al., 2013). High mortality typically follows the first infection by the fungi, also known as the killing front (Shigo, 1972). The beech scale does not appear to play a major role in the dispersal of *Neonectria*, but it is the most important factor predisposing trees to the disease (Cale et al., 2012; Lonsdale & Pratt, 1981). Other factors increasing disease susceptibility are the presence of birch margarodid (*Xylococculus betulae*) and low soil phosphorus availability (Cale et al., 2015).

The main BBD symptoms are bark cankers, scale colonies, and other bark lesions that are highly visible on the otherwise smooth bark. Physiologically, BBD makes it increasingly difficult for sap to flow, due to disruption of phloem and cambial tissues (Ehrlich, 1934). Thus, some later symptoms of BBD include leaf whitening, chlorosis, wilting, snagging, and eventually trunk breakage, due to water and nutrient limitation (Houston, 1994). These crown symptoms seem to be correlated with the presence of both beech scale and *Neonectria* fungi.

While mortality rates drop when the disease enters an endemic phase, the probability of secondary killing fronts in already-infected stands, also called “aftermath” stands, is expected to rise over time due to beech growth, suggesting a cyclic nature of BBD (Giencke et al., 2014; Houston, 1975). It is also expected that stand-level disease prevalence and severity will increase with climate change. A reduction in autumn precipitation, which usually causes detachment of beech scale from the bark during their mobile stage, and shorter cold seasons can favor scale insect population growth (Kasson & Livingston, 2012; Vincent et al., 2018).

At present, only sporadic monitoring efforts are carried out throughout the disease’s range. In Quebec, the Ministry of Natural Resources and Forests samples a relatively small number of trees each year with visual bark surveys (MFFP, 2020). Although the disease spreads relatively slowly, due to the immobile nature of the adult scale and to the fact that both insects and fungi must be present for an outbreak, an increased ability to detect BBD could assist in predicting new mortality phases in long-affected areas (e.g. eastern Quebec) or in newly affected areas such as eastern Ontario and Michigan (Cale et al., 2017).

Various remote sensing approaches have been used for the airborne identification of different plant diseases (Jackson, 1986; Klouček et al., 2019; Kothari et al., 2023; Senf et al., 2017; Weingarten et al., 2022). For example, oak wilt was detected with hyperspectral reflectance by Sapes et al. (2022). The use of spectroradiometers covering significant portions of visible and infrared light provides a non-invasive way to measure leaf structural and chemical properties (Asner & Martin, 2009, 2016; Kothari et al., 2023), which can be linked to disease symptoms. Such measurements capture leaf and canopy structure and composition, as these properties determine light reflection.

Hyperspectral data is defined as high-resolution spectral data covering the visible (380-700 nm), near infrared (NIR: 750-1000 nm) and short-wave infrared (SWIR: 1000-2500 nm) regions with hundreds of narrow bands. Reflectance variation for certain wavelengths, depending on their position in the spectrum, can be attributed to variation in the values of leaf and canopy functional traits (Asner & Martin, 2009). However, physiological stress was shown to be a more important determinant of spectral variation than specific functional traits or taxonomy, suggesting strong potential for hyperspectral data in pathogen detection (Fassnacht et al., 2022).

Extending the potential of hyperspectral data to a wider scale, beyond the leaf level, will require testing whether light reflectance at the canopy scale in airborne data can permit disease detection. However, the ability to predict canopy traits from leaf spectra has been highly variable, presumably due to major structural features of canopies. Canopy spectra are not only a function of the mean leaf spectra, but also the canopy’s biophysical properties (e.g., Leaf Area Index, LAI; Lead Angle Distribution, LAD), ground reflectance, sun exposure, and general canopy architecture (Asner, 1998; Goel, 1988; Jacquemoud et al., 1992; Jacquemoud & Ustin, 2019; Le Maire et al., 2004; Myneni et al., 1989; Ross, 1981). Airborne remotely sensed data are often summarized using derivative-based indices and spectral band ratios (e.g. NDVI), which can provide stronger discriminating capacity than leaf-level indices (Cho et al., 2008; Jacquemoud & Ustin, 2019; Le Maire et al., 2004). However, for several forest pathogens like oak wilt disease and Rapid ʻOhiʻa Death, PLSR has already been used successfully to detect symptoms at the canopy level (Sapes et al., 2022; Weingarten et al., 2022). Even though bark traits cannot be directly viewed with airborne data, making BBD detection a challenge, we hypothesized that disease advancement should influence leaf reflectance due to symptoms such as chlorosis, wilting, and the accumulation of phenolic compounds (Abdullah et al., 2018; Klouček et al., 2019).

At present, BBD monitoring is conducted at a small scale with mostly visual surveys of the bark (MFFP, 2020), and sparse spatial coverage. Here, we tested the ability of airborne hyperspectral imaging to detect BBD advancement. Very little is known about BBD impacts on leaf nutrient and water content, except for N:P ratios (Ouimet et al., 2015). Symptoms of BBD in the canopy are thought to appear in later stages of the infection (Houston, 1994) and we hypothesized that the main regions of interest in the spectra would be those related to water content (SWIR) and to pigment concentration in the visible region. Structural symptoms in the SWIR might also be of some importance given that leaf loss is a BBD symptom (Ehrlich, 1934). Hence, for diseased individuals, we expected lower reflectance values in the green section of the spectra due to reduced chlorophyll, and lower reflectance values overall due to leaf loss and canopy damage. We also assessed the potential of close-range leaf-level hyperspectral data, which eliminates atmospheric interference, to predict BBD progression. Secondarily, we tested for multivariate correlations between hyperspectral signatures at canopy and leaf level to assess the potential transferability of models across scales.

## 2. Methods

### 2.1 Study area

This study took place in Mont-Saint-Bruno National Park, an 8.9 km^2^ protected area in southern Quebec, Canada, 15 km east of Montréal (Figure 1a). The area is in the Saint Lawrence Lowlands’ bitternut hickory - maple bioclimatic domain, with an elevation of 48-218 m above sea level. The site has a high abundance of beech, and most stands in the region are already affected by BBD, still experiencing high mortality (MFFP, 2020). Total annual precipitation is roughly 1000 mm; the average temperature is 6.3 °C.

**Fig. 1.**
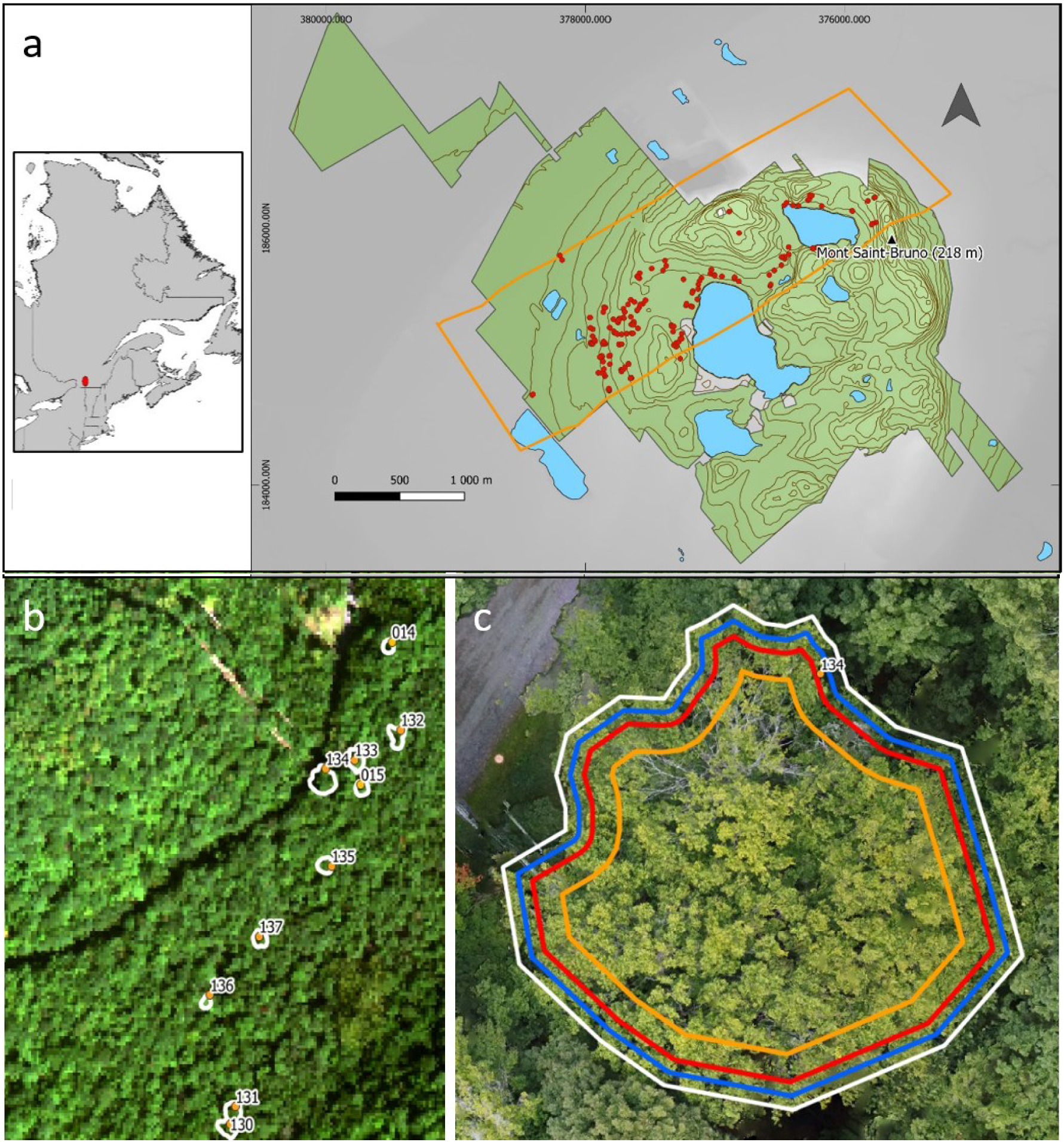
Overlaying of the airborne hyperspectral data and the drone-based data, enabling the precise extraction of pixels corresponding to a selected individual’s canopy. Study area (the flight line is shown by the yellow polygon and the sampled beech trees by the red dots). b. A portion of the hyperspectral imagery and delineated beech canopies. c. Canopy buffers (blue = 0.5 m, red = 1 m, yellow = 2 m), shown for one delineated canopy.

### 2.2 Data acquisition

#### Airborne data

The airborne hyperspectral survey was conducted on September 8^th^, 2018, using a Twin Otter fixed wing aircraft operated by the National Research Council of Canada’s Flight Research Lab. A pushbroom imager, with a field of view angle of 39.8°, compiled data from two hyperspectral sensors (CASI, 288 bands from 375 to 1061 nm, and SASI, 100 bands from 957 to 2442 nm). One flight line covered the study area with a resolution of 2 m × 2 m, thus eliminating potential variation between flight lines (Figure 2.1a). Radiometric and atmospheric corrections based on the ATCOR-4 model were executed on the raw hyperspectral data as standard pre-processing steps (Inamdar et al., 2021). All airborne hyperspectral data were drawn from the CABO database (Canadian Airborne Biodiversity Observatory).

**Fig. 2.**
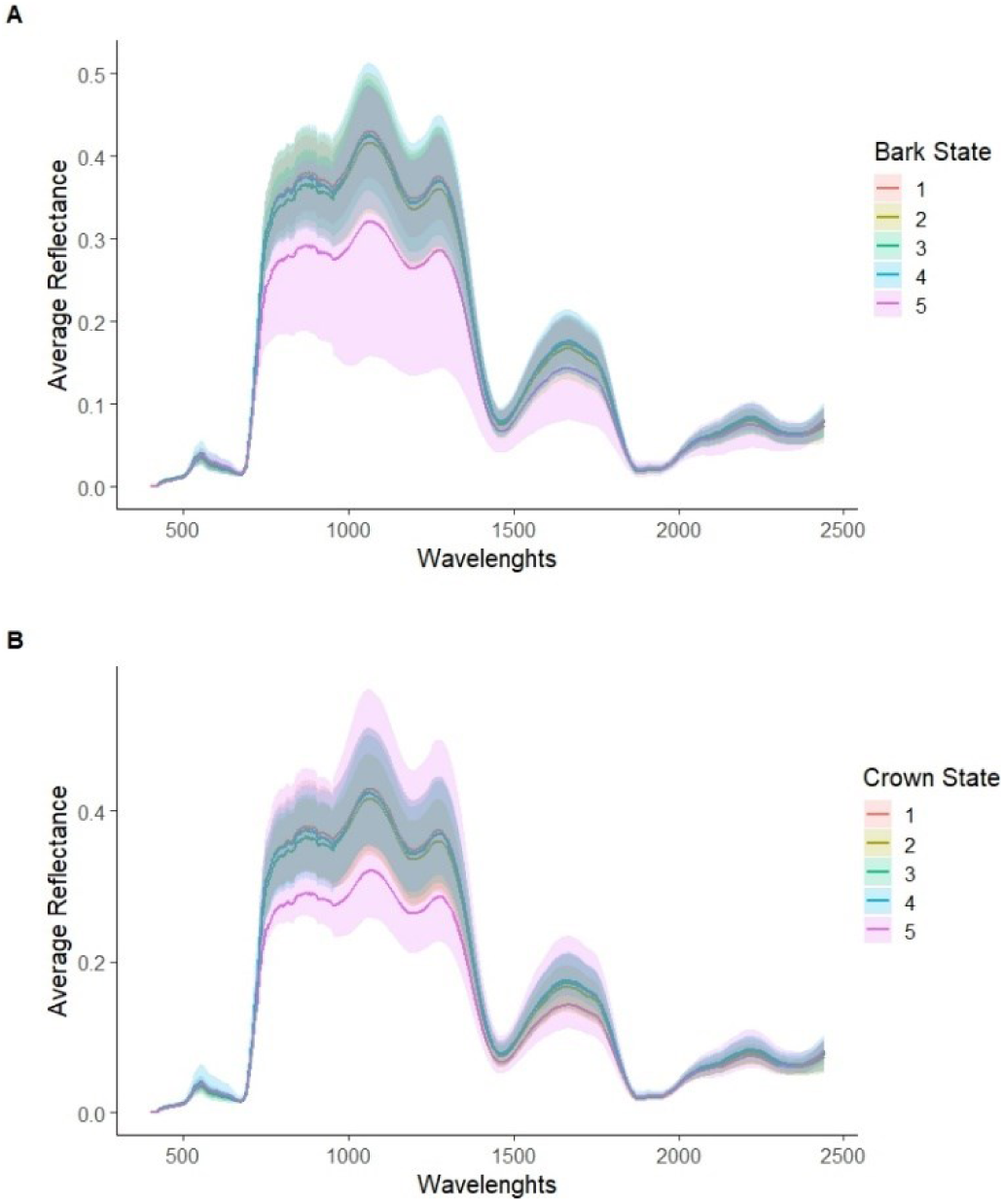
Average and range of reflectance for trees with a variety of scores for bark state and crown state (input variables for our disease severity score). These reflectance spectra correspond to beech trees sampled at the canopy level.

The airborne hyperspectral imagery was aligned and spatially linked with the canopies of the sampled beech trees in the field. However, due to the relatively coarse spatial resolution (∼2 m) of the hyperspectral imagery, we were unable to confidently delineate canopies using only these data. As such, we also gathered drone-based red, green and blue (RGB) imagery above the sampled beech trees. Photographs of the canopy taken with a photogrammetric configuration provided a detailed visual account of the site. Specifically, a consumer Unoccupied Aerial Vehicle (UAV) constructed by DJI, the Mini 2, was used for flights covering 18 portions of the study site (Figure S1). These regions were used as flight polygons corresponding to approximately 0.63 km^2^ in total. The flight pattern involved back-and-forth movement with 90% frontal overlap and 70% lateral overlap between photos, taken at a mean height of 40 m above ground level (AGL), perpendicular to the ground. We overlaid drone and plane-based imagery, permitting precise beech canopy positioning and crown delineation; this method reduces uncertainty in the georectification of hyperspectral imaging even with few ground control points (GCPs) (Habib et al., 2016). To maintain close-range resolution with < 1 m horizontal and vertical error, at least three GCPs were positioned evenly across every drone flight polygon. GCP locations were measured with < 30 cm error using a Trimble Catalyst device with RTK positioning. Some GCPs were difficult to locate through the forest canopy, so during post-processing we added some control points (with a lower precision) based on objects visible from above, such as dead wood or canopy gaps.

#### Beech Tree Measurements

We measured disease-related factors on 160 beech trees distributed as evenly as possible throughout the park in summer 2022. We estimated four proxies of disease severity: bark state, canopy integrity, perithecia cover, and scale insect cover. For all four variables, a tree (or portions of a tree) was assigned a categorical score between 1 and 5.

Bark state was assessed based on the percentage of cankers/injuries covering each of four sides of a tree, one side in each cardinal direction (N, S, E, W). (1) 0-20% cover of cankers/injury, (2) 20-40%, (3) 40-60%, (4) 60-80% and (5) 80-100%. A tree face with no bark was scored as a 5.

For canopy integrity, defoliation percentages defined five disease classes: (1) no apparent crown damage due to BBD; (2) < 25% defoliation, (3) 25-50%, (4) 50-75%, and (5) 75-100%. For trees with no canopy (dead or alive) due to trunk snapping, canopy integrity was scored as a 5.

For perithecia cover, we used the following semi-quantitative scale: (1) No perithecia visible for the present year or recent years; (2) Very few perithecia visible on the bark; (3) Some visible signs of perithecia but no red globose bodies present; (4) Presence of *Neonectria* red fruiting bodies and significant signs of perithecia; (5) High cover of *Neonectria* fruiting bodies.

For scale cover, we used the following categories: (1) No visible scale; (2) 1 or 2 small scale colonies; (3) more than 2 colonies but only small ones; (4) a moderate density of larger scale colonies; and (5) a high density of larger colonies, often almost covering the whole bark. For this step, *Cryptococcus fagisuga* and *Xylococculus betulae* were both considered as scale insects and were not differentiated. Note that neither the presence of the beech scale nor that of the *Xylococculus* insect was precisely quantified, as they do not directly affect the tree’s vascular function and therefore should not cause foliar symptoms.

Finally, every beech canopy was also positioned spatially with a GPS using GNSS corrections allowing < 1 m precision.

#### Leaf-level spectroscopy

In summer 2023, we collected leaf samples on 37 individuals out of the original 160. We selected individuals to represent a wide range of disease levels (approximately as many healthy individuals as severely affected individuals and a gradient of infection intensity). Leaf sampling was conducted between August 31^st^ and September 14^th^ to match the phenological stage of the canopies in the airborne imagery. Leaf samples were harvested using a Big Shot arborist catapult (Sherrilltree Inc., Greensboro, North Carolina, USA) with launch weights (12-20 oz) and an ultralight nylon wire ∼ 75 m long (Youngentob et al., 2016).

After collection, leaves were stored in inflated sealable plastic bags placed in a cooler for a maximum of 5 h; leaves were kept on their branches to allow water and nutrients to circulate. We then obtained leaf-level spectroscopy data on-site, using the HR-1024i field spectroradiometer from Spectra Vista Corporation (Poughkeepsie, New York, USA) combined with their 3-inch Reflectance/Transmission Sphere (Laliberté & Soffer, 2018). These data covered wavelengths from 340 to 2500 nm (2161 spectral bands).

### 2.3 Data processing

We aligned drone canopy photos using Agisoft Metashape, version 2.0.2.16404, which conducts photogrammetric processing for series of digital images (Agisoft, 2019). For each flight polygon, photos were aligned and positioned in space using the GCPs as spatial references.

Delineation of hyperspectral pixels within the canopies of our field-sampled trees was done using QGIS, version 3.16 (QGIS Development Team, 2023). We first overlaid the airborne hyperspectral data with the drone-based imagery. Canopy polygons that could be identified with an error of ≤ 1 m horizontally were drawn manually from the drone imagery (Figure 1b). This was possible for 126 out of 160 trees, such that 34 individuals were excluded from further analysis. We then used R (version 4.2.3) to extract the hyperspectral pixels falling within the manually drawn polygons using the “terra” and “exactextractr” packages (Baston, 2022; Hijmans, 2023).

Spectral data were combined across the CASI and SASI sensors. For every overlapping wavelength (between 957.5 and 1061.5 nm), we matched reflectance values from both sensors using the “spectrolab” package (Meireles et al., 2017). Using the “hsdar” package (Lehnert et al., 2019), we then implemented several processing steps to reduce the noise generated by the interaction of light with the atmosphere. Specifically, we consecutively performed a masking of the atmospheric absorption regions (from 350 to 400 nm and from 920 to 957 nm), an interpolation of reflectance in these regions, and an application of the Savitzky-Golay smoothing filter with a size window of 11 bands and a polynomial of order 2 (Savitzky & Golay, 1964).

Given some uncertainty in canopy delineation, we implemented a spatial buffer to eliminate pixels at the canopy edge. On the 126 canopies, we tested buffer sizes of 0.5 m, 1 m, and 2 m, eliminating hyperspectral data closer to the canopy edge than these cutoffs (Figure 1c). A pixel was included if more than 50% of it fell within the buffer (pixel weight ≥ 0.5). For some buffers, some canopies no longer contained any pixels, resulting in sample sizes of n=126 (0.5 m buffer), n=122 (1 m) and n=80 (2 m). Finally, we filtered shadowed pixels by eliminating those with reflectance <0.0075 at the 679.11 nm band, which corresponds to the red color band that was shown to be the best measure of shadow evaluation for this site (Wallis et al., 2023).

### 2.4 Statistical analysis

To characterize disease severity, we used only the bark state and canopy state variables. Beech scale cover appeared to vary haphazardly with foliar symptoms observed in the field, and in a previous study, beech scale abundance was not linked to other disease symptoms (Cale et al., 2015; Teale et al., 2009). We found a moderate correlation (r=0.53) between the canopy state and bark state variables and a strong correlation between perithecia cover and bark state (r=0.83); perithecia cover was thus considered redundant. To generate one continuous disease severity variable, we used the main axis of a principal components analysis (PCA) with only bark state and canopy state; PC1 explained 73.4% of the variance in these two variables.

Partial least-squared regression (PLSR) models were performed using the airborne hyperspectral data to predict the composite disease severity variable using the same settings as Burnett et al. (2021) and the same validation scheme as Kothari et al. (2023). A separate model was run for each of the three buffer distances; each model considered the whole reflectance spectrum as a predictor. For these analyses, we used the “spectratrait” package in R developed by Burnett et al. (2021), implementing a Jackknife internal cross-validation on separate training and validation data subsets with ∼80% and ∼20% of the dataset, respectively. We further divided the calibration set randomly into subsets, with approximately 70% for training and 30% for validation. This division was repeated 100 times to evaluate the predictive models’ performance and reliability, as described by Kothari et al. (2023). This procedure allowed us to generate distributions of key statistics (R^2^ and RMSE) that assess variation in model performance in response to random variations in both the training and test datasets. %RMSE was calculated by dividing the RMSE by 95% of the range in severity score values, serving as a measure of robustness to outliers (Kothari et al., 2023). The number of components was selected for each model with the “FirstPlateau” method from the same package. For our best model, with the use of Jackknife regression coefficients (obtained by refitting the model after leaving one observation out each time, then averaging the predictors’ coefficients), we determined the magnitude and sign of each wavelength’s and spectral region’s contribution to disease score prediction. Coefficient values near 0 signify a weak contribution to prediction. Finally, variable influence on projection (VIP) scores were calculated in order to assess the relative importance of each wavelength in explaining disease severity (Burnett et al., 2021). VIP scores > 0.8 indicate wavelengths with a high importance in prediction (Wold et al., 2001). We therefore conducted a subsequent PLSR model only using the spectral bands with VIP > 0.8 as predictors.

We checked for patterns of spatial autocorrelation in the hyperspectral data across trees using the “gstat” package (Pebesma, 2004); we found no convergence of the semi-variance in these models, indicating no spatial autocorrelation. For the analysis of leaf-level hyperspectral data, PLSR models were run in the same way as for airborne data.

To test the correlation between airborne and leaf hyperspectral data, we performed a Procrustes analysis using the 29 sampled trees common to the airborne (buffer = 0.5 m) and leaf-level data, using the “vegan” package (Oksanen et al., 2022).

Finally, we tested whether some widely used remotely sensed vegetation and water indices correlated with disease progression (Table 1). Because PLSR models are based on linear combinations of predictors, they will not necessarily capture information in spectral indices that are based on ratios of wavelengths (or ratios of sums or differences). Specifically, we analyzed vegetation difference (greenness) and water content indices, which we might expect to change during BBD progression (Cale et al., 2015; Ouimet et al., 2015). Specifically, the DD index (Double Difference), NDVI (Normalized Difference Vegetation Index), gNDVI (Green NDVI), NPQI (Normalized Phaeophytinization Index) and REIP (Red-edge Inflexion Point) capture greenness in different ways, and NDWI (Normalized Difference Water Index) captures canopy water content. Multiple studies have demonstrated that NDVI provides a useful assessment of vegetation health and stress (Dash et al., 2017; Guerra-Hernández et al., 2021; Verbesselt et al., 2009). NPQI can be of some importance in crown health assessment (Pontius et al., 2008). Simple linear regression models were used to assess these relationships. AIC was used for model selection (Akaike, 1973). All statistical analysis were carried out on R (version 4.2.3; R Core Team, 2021).

**Table 1.** Description of the water content and vegetation indices used as predictors for BBD progression in an exploratory approach.

| *Index* | *Leaf trait related* | *Formula* | *Reference* |
| --- | --- | --- | --- |
| NDWI | EWT and LMA | $\frac{R_{860nm} - R_{1240nm}}{R_{860nm} + R_{1240nm}}$ | (Gao, 1996) |
| NDVI | Chl + LAI | $\frac{\overline{R_{NIR; 765:815nm}} - \overline{R_{Red; 636:686nm}}}{\overline{R_{NIR; 765:815nm}} + \overline{R_{Red; 636:686nm}}}$ | (Rouse Jr. et al., 1973) |
| gNDVI | Chl a | $\frac{R_{750nm} - \overline{R_{Green; 526:576nm}}}{R_{750nm} + \overline{R_{Green; 526:576nm}}}$ | (Gitelson et al., 1996) |
| NPQI | Stress & Chl | $\frac{R_{415nm} - R_{435nm}}{R_{415nm} + R_{435nm}}$ | (Barnes et al., 1992) |
| REIP | Chl & Chl x LAI | $R''(\lambda_i) = 0$ | (Horler et al., 1983) |
| DD | Chl | $(R_{749nm} - R_{720nm}) - (R_{701nm} - R_{672nm})$ | (Le Maire et al., 2004) |

$R_\lambda$ = Reflectance at a wavelength $\lambda$ ; EWT = Equivalent Water Thickness; LMA = Leaf Mass Area; Chl = Chlorophyll; LAI = Leaf Area Index. Related leaf traits were obtained from Le Maire et al. (2004).

## 3. Results

We found minimal correspondence between the hyperspectral signatures of trees and our two disease predictors (bark state and crown state) (Figure 2). Only the most advanced stages of infection (bark state = 5 and crown state = 5) showed visually detectable differences in reflectance variation at the canopy level.

### Full-spectra models

Overall, airborne hyperspectral data had a limited capacity to predict beech bark disease severity, regardless of the three buffers used for canopy pixel selection. Disease score values (response variable) ranged between −1.58 and 3.26, with higher values representing more severe infection. Each model showed different 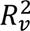 (validation) values going as high as 0.24, but as low as 0. The model with a buffer of 2 m had the highest *R*^2^in validation analyses, equal to 0.24, and a % RMSE of 30.2% (Table 2). However, due to its lower 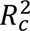 (calibration) value (0.13) – indicative of underfitting and model instability – and to the uncertainty in 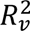 with the 2 m buffer (ranging between 0.02 and 0.37; Figure 3), we selected the 0.5 m buffer model to conduct further analyses (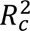=0.18 and 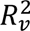=0.10).

**Fig. 3.**
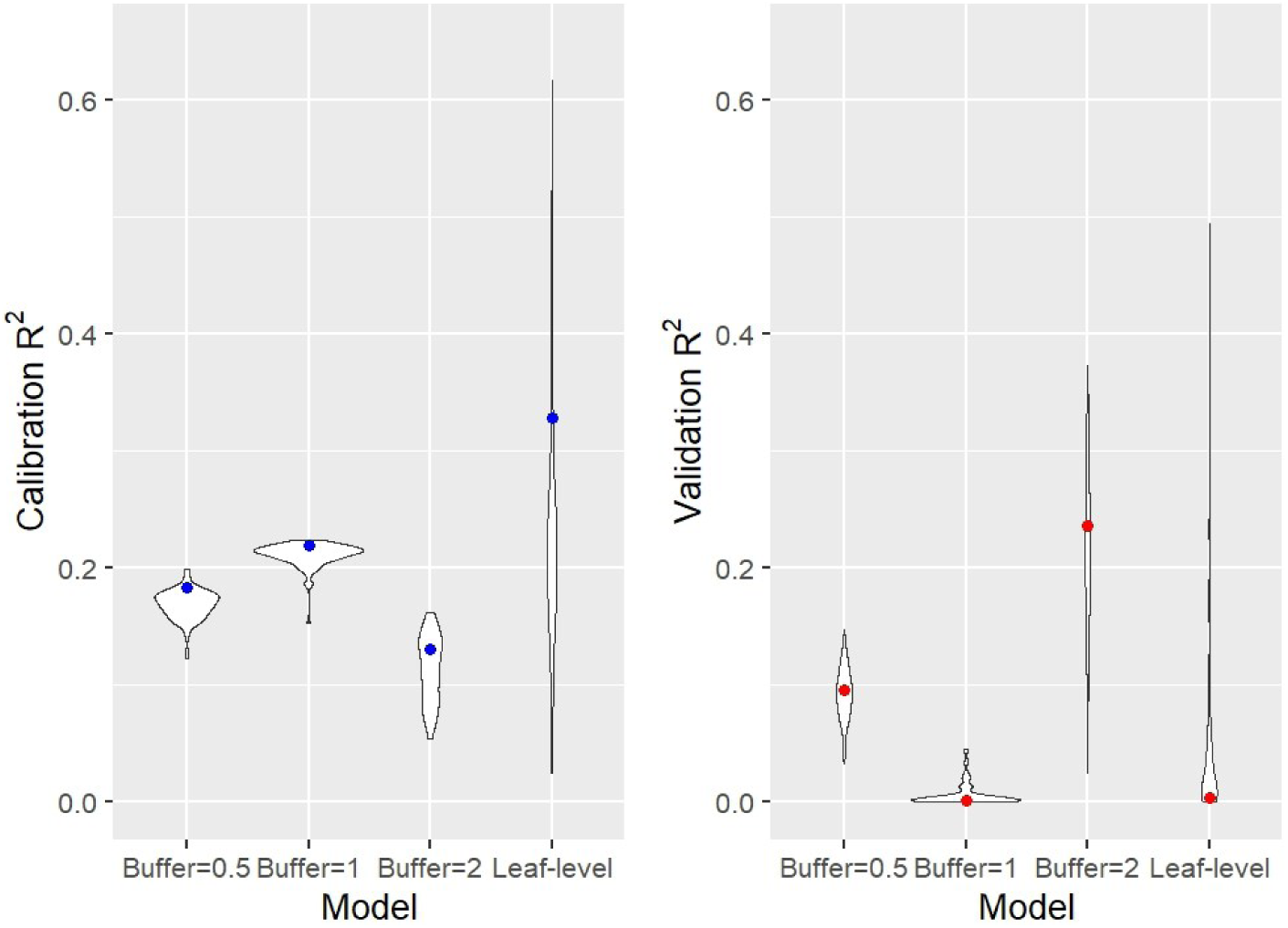
Violin plots showing the range of R^2^ values for both calibration (blue, left) and validation (red, right) full-spectra PLSR models. The dot is the averaged R^2^ value in each performed model.

**Table 2.** PLSR model results based on the airborne and the leaf-level hyperspectral datasets.

| Buffer | NoC | $R_c^2$ | $RMSE_c$ | $R_v^2$ | $RMSE_v$ | % $RMSE$ |
| --- | --- | --- | --- | --- | --- | --- |
| *Airborne models* |  |  |  |  |  |  |
| 0.5 m | 3 | 0.18 | 0.98 | 0.10 | 1.29 | 32.5 |
| 1 m | 3 | 0.22 | 1.02 | 0.00 | 1.16 | 32.3 |
| 2 m | 2 | 0.13 | 1.14 | 0.24 | 1.37 | 30.2 |
| *Leaf-level model* |  |  |  |  |  |  |
|  | 3 | 0.33 | 0.81 | 0.00 | 0.91 | 40.0 |

NoC is the Number of Components used in each PLSR model, $R_c^2$ and $R_v^2$ are the explained variance proportions for the calibration and the validation models, respectively, and $RMSE_c$ and $RMSE_v$ are the root mean squared of error predicted as a disease score. %RMSE thus refers to prediction variation compared to total disease score range.

For the model with a 0.5 m buffer, VIP scores peaked in the green-yellow part of the visible region (∼560 nm), in the red-edge region (∼720 nm) and in the SWIR region at ∼1300 nm, ∼1675 nm and ∼2250 nm, sometimes reaching scores >2 (Figure 4). The jackknife validation coefficients confirmed the relative importance of the predictors, particularly for the red-edge region with a positive relative contribution on disease score of 0.90 (Figure S2).

**Fig. 4.**
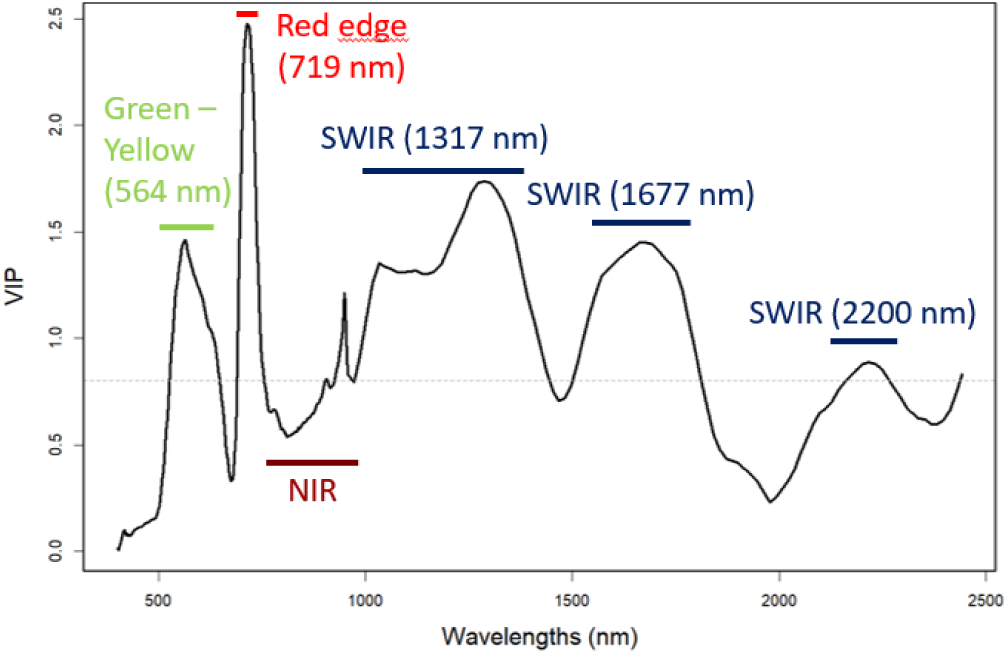
Variable influence on projection (VIP) scores across wavelengths for the best airborne hyperspectral model (buffer=0.5 m). Regions > 0.8 (above the dashed grey line) are considered to have high importance. Green-Yellow bands are situated in the visible region; Red-edge is the region between the visible and NIR. For each region, the wavelength with the maximum score was indicated, except for the NIR region, which did not make a significant contribution to our model’s predictive capabilities.

At the leaf-level, the PLSR model gave, for calibration, an 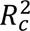 of 0.33 with an RMSE of 0.81. For validation, 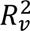 was 0 with a %RMSE of 40.0% (Table 2). Due to the poor performance of the leaf-level PLSR model, VIP scores were not considered meaningful for understanding the results. The jackknife validation coefficients for this model showed that the contribution to disease severity was very low for the whole spectra, only reaching maximum contribution to disease score of ∼0.06, also in the red-edge region (Figure S3).

The symmetric Procrustes analysis comparing the airborne and leaf-level datasets resulted in a correlation of r = 0.326, although this was not statistically significant (p = 0.106).

### VIP-only spectral model

Another PLSR model was run using only bands with VIP scores >0.8 from the best full-spectra model (model with 0.5 m buffer). This model used 132 bands as predictors, situated between 537-750 nm; 957-1812 nm and 2127-2442 nm. This model resulted in 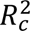 of 0.12 and 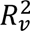 of 0.48 with a %RMSE of 27.2% (NoC = 2; RMSE_c_ =1.19; RMSE_v_ = 1.07).

### Spectral index models

Using the same pixel selection criteria as for other analyses, we found significant relationships between disease progression and different band ratios. The band ratios used were based on wavelengths showing VIP scores >0.8. The best model using band ratios was the one combining gNDVI, NPQI and REIP (R^2^=0.19; ΔAIC=23), indicating an influence of greenness, canopy density and leaf stress in predicting BBD (Table 3). For most models, greenness indices explained significant portions of the variation in BBD symptoms, while this was not the case for NDWI.

**Table 3.** Linear models predicting BBD progression from band ratios.

| *Models* | *Predictors* | *p-values* | *R<sup>2</sup>* | *ΔAIC* |
| --- | --- | --- | --- | --- |
| Intercept-only | ~1 | - | - | 0 |
| Water content index | ~NDWI | 0.861 | -0.01 | -2 |
| Vegetation indices | ~DD*** | 5.88E-04 | 0.08 | 10 |
|  | ~NDVI*** | 2.71E-04 | 0.09 | 11 |
|  | ~gNDVI*** | 1.09E-04 | 0.11 | 13 |
|  | ~NPQI* | 0.023 | 0.03 | 3 |
|  | ~REIP | 0.054 | 0.02 | 2 |
|  | ~gNDVI***+NPQI***+REIP* | 3.28E-06;<br>2.67E-04;<br>0.018 | 0.19 | 23 |
| Water and vegetation indices | ~NDWI+DD*** | 0.161;<br>2.20E-04 | 0.09 | 10 |
|  | ~NDWI+NDVI*** | 0.236;<br>1.38E-04 | 0.10 | 11 |
|  | ~NDWI+gNDVI*** | 0.054;<br>1.65E-05 | 0.12 | 15 |

Significance thresholds: \*: $p<0.05$ , \*\*: $p<0.01$ , \*\*\*: $p<0.001$

## 4. Discussion

Our results revealed weak predictive relationships between hyperspectral data and beech bark disease severity, in both canopy-level and leaf-level models. This suggests that changes in canopy or leaf traits during most of disease progression are either absent or undetectable, except for already known symptoms in the most severe cases of infection, such as chlorosis (a decrease in chlorophyll pigments) and changes in water content. Even the best predictive models of BBD severity (using spectral indices) accounted for only 19% of the total variation in disease severity at a spatial resolution of 2 m × 2 m.

Poor model performance could potentially be explained by the disease’s vascular nature. As necrotic phloem is not uniformly distributed throughout the trunk, sap flow might not be altered at a consistent time during disease progression or in the same way in all parts of the tree (Ehrlich, 1934; Houston, 1994). If there is temporal randomness in the onset of leaf or canopy symptoms, this would diminish our model’s ability to predict disease progression. In addition, if symptoms occur only in small portions of the crown, accuracy of disease assessment would be limited with an image resolution as high as 2 m × 2 m. For infections reaching the canopy, where symptoms appear at branch level, lateral strips link affected trunk sections to diseased branches. (Shigo, 1962). Indeed, BBD symptoms could be diluted among parts of the canopy, making them appear healthy due to the large pixel size (relative to leaf and branch dimensions) and a small number of pixels per canopy. Consistent with this possibility, our best airborne PLSR model conserved the most pixels per canopy (buffer=0.5 m), although it is possible that the 2 m buffer model (n=80) would have performed better with a larger number of tree crowns (Goodhue et al., 2012). Our VIP-only PLSR model was not interpretable, given high dissimilarity between the results of the calibration and validation models, and unstable R^2^ values (Figure S4). Localized canopy damage might also help explain the failure of our leaf-level model, since highly damaged branches might have no leaves, and sampling one branch would not represent disease severity of the whole tree.

Even if minimal variance in disease severity was explained by airborne models (R^2^=0.10), the results revealed some information about canopy symptoms of beech bark disease. For our best model, VIP scores peaked in the green-yellow part of the visible region, in the red-edge region and in a significant part of the SWIR region, suggesting a link between disease severity and chlorophyll content, water content, and leaf dry matter content (Curran et al., 2001; Jacquemoud & Ustin, 2019; Kokaly & Clark, 1999). Further support for the importance of chlorophyll (chlorophyll a specifically), LAI and leaf stress came from our best spectral indices’ models, for which three vegetation indices commonly used for their assessment came out significant (gNDVI, NPQI and REIP; R^2^=0.19). Since traits are often linked to multiple bands simultaneously, differential indices such as gNDVI are more sensitive than unique bands to water and nutrient variation (Le Maire et al., 2004; Weingarten et al., 2022). Ratio-based indices likely provide a better assessment of chlorophyll differences in the canopy, as they are more sensitive to a wider range of chlorophyll concentrations (Gamon & Surfus, 1999; Jacquemoud & Ustin, 2019).

Our Procrustes analysis showed no relationship between airborne and leaf-level hyperspectral measurements. This suggests that canopy measurements are influenced not only by leaf characteristics, but perhaps even more strongly by canopy structure and lighting conditions during data acquisition (Asner, 1998; Myneni et al., 1989; Ross, 1981). These biophysical constraints have a major impact on reflectance at the canopy level, suggesting that spectral data at the leaf level cannot be easily scaled up to airborne canopy measurements, at least within species, as in our study.

Overall, using current best practices, we did not find disease severity to be predictable using spectral data. As suggested by Lopez et al. (2023), this disease might be more readily diagnosed using structural and textural features instead of spectral ones. In our study, the most important canopy symptoms, which could only be detected in later stages of infection after fungi have already caused vascular damage, corresponded to classic canopy symptoms of beech bark disease (Ehrlich (1934): canopy yellowing and leaf/branch loss. Except for spectral features corresponding to pigment content and water absorption, no other spectral regions were indicative of early symptoms that could help with disease prediction before it causes heavy damage. Thus, we would not recommend hyperspectral data for beech bark disease severity assessment, particularly in mildly and moderately affected stands. It remains possible that RGB imaging and LiDAR data, combined with machine learning and artificial intelligence approaches, may help improve predictions of severity for bark diseases, by focusing on canopy structure at a finer resolution (Lopez et al., 2023; Marvasti-Zadeh et al., 2023). Considering intra-canopy variability in BBD-affected trees, such methods could lead to more accurate predictions of BBD severity progression in American Northeastern forests.

## Acknowledgements

This study was conducted in the framework of CABO (the Canadian Airborne Biodiversity Observatory), funded by the Natural Sciences and Engineering Research Council of Canada (RGPIN-509190-2017). Therefore, we thank NSERC, as well as all those involved in acquiring hyperspectral data at the Saint-Bruno study site (Pablo Arroyo-Mora among others), and the team that processed the hyperspectral imagery, including Deep Inamdar and Margaret Kalacska. We would also like to thank the administration of Mont-Saint-Bruno National Park for their collaboration and ongoing assistance with the acquisition of permits and authorizations. We thank Audrey Thériault and Léa Gauthier-Soumis for their much-appreciated help as field assistants. Finally, we are grateful to the members of the Vellend laboratory for their valuable advice and support.

## Abbreviations

AGL: Above Ground-Level

BBD: Beech Bark Disease

CABO: Canadian Airborne Biodiversity Observatory

GCP: Ground control point

GIS: Geographic Information System

GNSS: Global Navigation Satellite System

HS: Hyperspectral

NIR: Near Infrared

PCA: Principal Components Analysis

PLSR: Partial least-squared regression

PPK: Post-Processed Kinematic GNSS data

RGB: Red-Green-Blue

RMSEP: Root Mean Square Error Predicted

RTK: Real-Time Kinematic GNSS data

SWIR: Short-Wave Infrared

UAV: Unmanned Aerial Vehicle

VIP: Variable Importance in Projection

VIS: Visible light

## Notes

### Competing Interest Statement

The authors have declared no competing interest.

