## Supplemental Figures (S1 to S4) for "Hyperspectral imaging has a limited ability to remotely sense the onset of beech bark disease"

[1]: Université de Sherbrooke, Department of Biology

[2]: Université de Montréal, Department of Biological Sciences

[3] : Technische Universitat Berlin

### Supplementary Materials

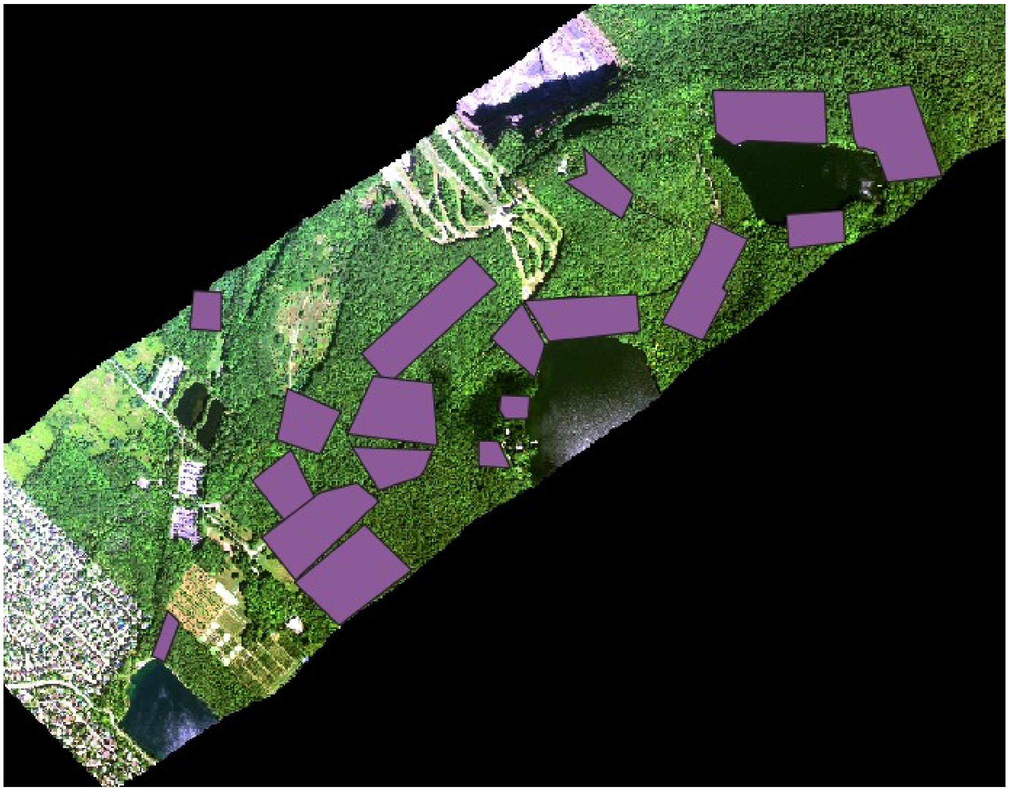

**Figure S1.** **RGB drone imagery flight polygons (18 different flight zones, covering a total area of ~63 hectares).**

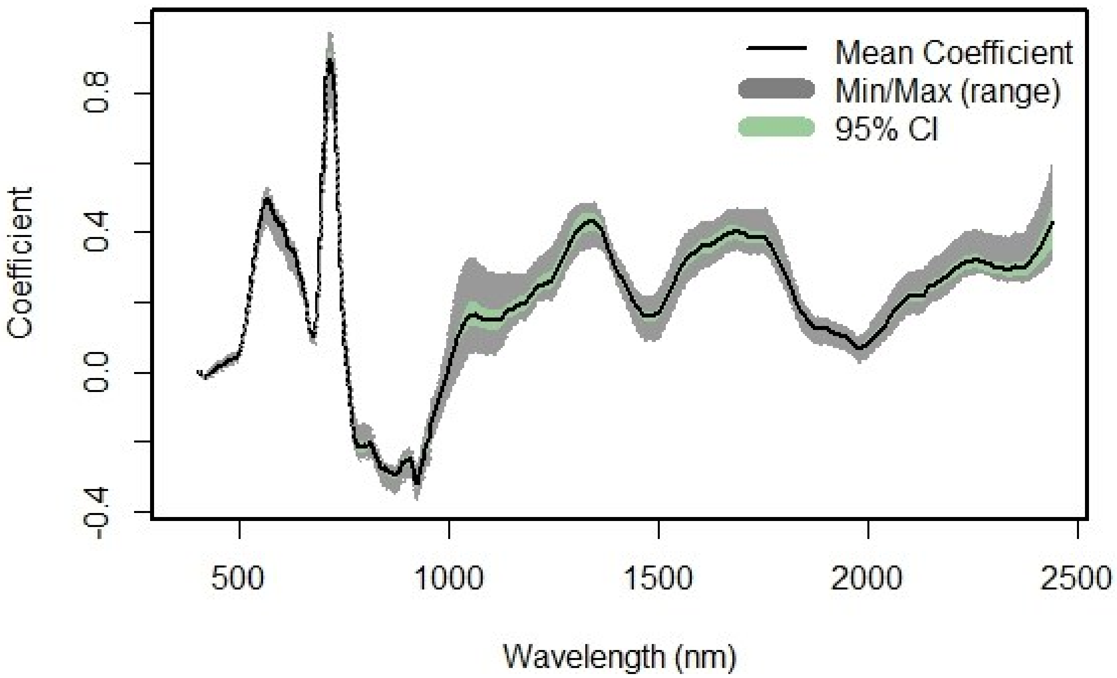


**Figure S2. Jackknife validation results for model coefficients in the best airborne PLSR model (buffer = 0.5 m).**

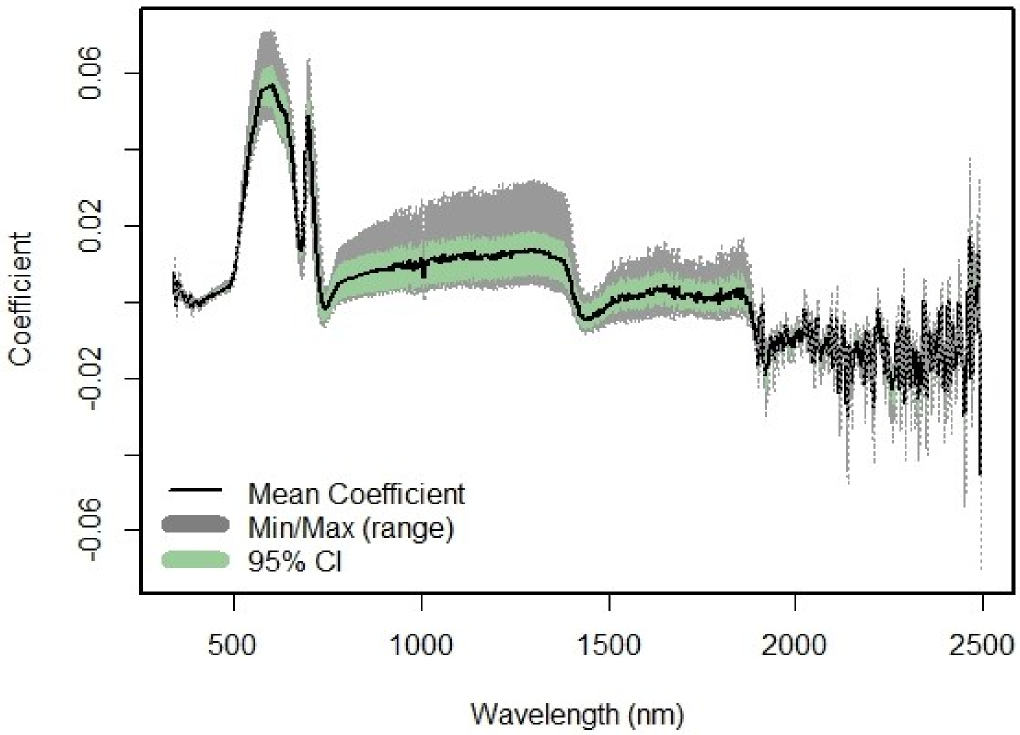


**Figure S3. Jackknife validation results for model coefficients in the best leaf-level PLSR model.**

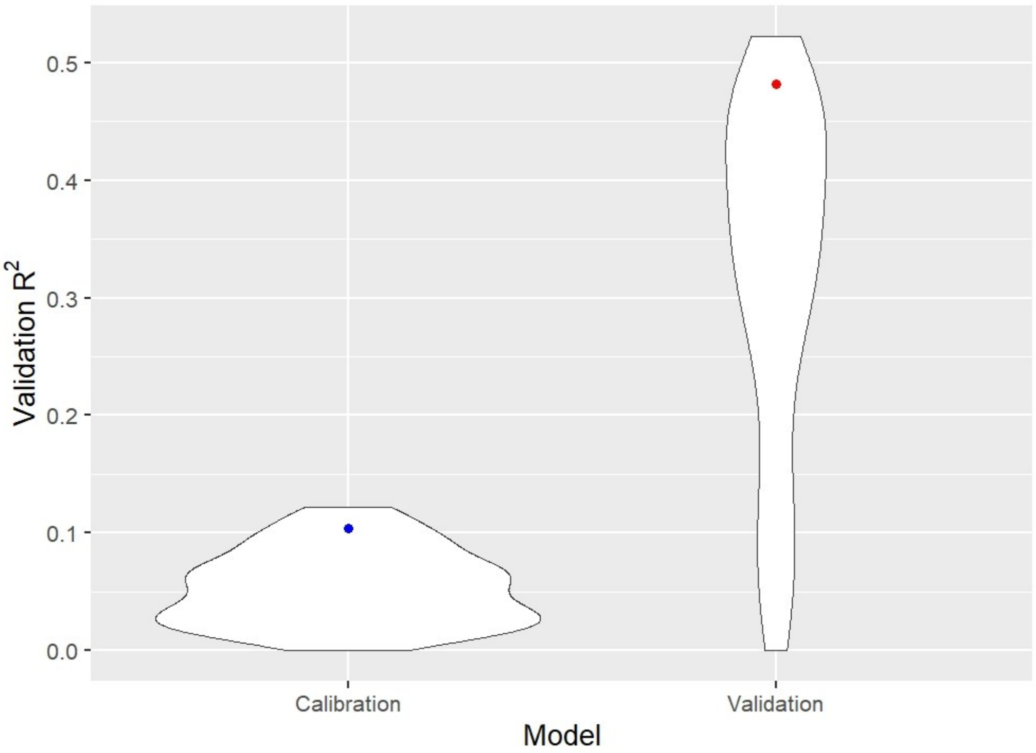


**Figure S4. Violin plots showing the range of $R^2$ values for both calibration (blue, left) and validation (red, right) VIP-only PLSR models.** The dot is the averaged $R^2$ value in each performed model.
